# Brain activation during the N-back working memory task in individuals with spinal cord injury: a functional near-infrared spectroscopy study

**DOI:** 10.1101/2024.02.09.579655

**Authors:** Donna Y. Chen, Xin Di, Nayyar Amaya, Hai Sun, Saikat Pal, Bharat B. Biswal

**Author notes:** Correspondence: Bharat B. Biswal, Ph.D., 607 Fenster Hall, University Heights, Newark, NJ, 07102, US.

## Abstract

Cognitive impairments have frequently been reported in individuals with spinal cord injury (SCI) across different domains such as working memory, attention, and executive function. The mechanism of cognitive impairment after SCI is not well understood due to the heterogeneity of SCI sample populations, and may possibly be due to factors such as cardiovascular dysfunction, concomitant traumatic brain injury (TBI), hypoxia, sleep disorders, and body temperature dysregulation. In this study, we implement the Neuropsychiatric Unit Cognitive Assessment Tool (NUCOG) to assess cognitive differences between individuals with SCI and age-matched able-bodied (AB) controls. We then use an N-back working memory task and functional near-infrared spectroscopy (fNIRS) to elucidate the neurovascular correlates of cognitive function in individuals with SCI. We observed significant differences between the SCI and AB groups on measures of executive function on the NUCOG test. On the N-back task, across the three levels of difficulty: 0-back, 2-back, and 3-back, no significant differences were observed between the SCI and AB group; however, both groups performed worse as the level of difficulty increased. Although there were no significant differences in N-back performance scores between the two groups, functional brain hemodynamic activity differences were observed between the SCI and AB groups, with the SCI group exhibiting higher maximum oxygenated hemoglobin concentration in the right inferior parietal lobe. These findings support the use of fNIRS to study cognitive function in individuals with SCI and may provide a useful tool during rehabilitation to obtain quantitative functional brain activity metrics.

## 1. Introduction

Every year in the U.S., there are approximately 18,000 new cases of traumatic spinal cord injury (SCI), as of 2022 (National Spinal Cord Injury Statistical Center, Birmingham, AL, 2022). Cases of traumatic SCI are caused primarily by events such as vehicle accidents, falls, acts of violence, and sports/recreational activities. After injury, individuals with SCI are at a 13 times greater risk of cognitive impairment than able-bodied individuals, in addition to being at a high risk for developing pervasive mental disorders such as depression and anxiety (Craig et al., 2017; Lim et al., 2017; Pasipanodya et al., 2021; Pozzato et al., 2023). Furthermore, individuals with SCI typically score lower on cognitive tests evaluating information processing speed, verbal learning and memory, and verbal fluency than age-matched able-bodied individuals (Chiaravalloti et al., 2018). The mechanism of cognitive decline after SCI is not well understood, with studies suggesting cardiovascular dysfunction (Nightingale et al., 2020; Wecht & Bauman, 2013), concomitant traumatic brain injury (TBI) (Bradbury et al., 2008; Macciocchi et al., 2013; Nott et al., 2014), hypoxia (Hernandez-Gerez et al., 2019), sleep disorders such as obstructive sleep apnea (Carlozzi et al., 2022; Pasipanodya et al., 2021; Shafazand et al., 2019), and body temperature dysregulation (Wecht et al., 2015) as possible causes of cognitive impairment in individuals with SCI (Alcántar-Garibay et al., 2022). Despite evidence of cognitive decline in individuals with SCI, the extent to which cognitive decline alters functional brain reorganization is unclear. This is partially attributed to the lack of neuroimaging studies focused on understanding cognitive function after SCI.

Structural and functional brain changes have been observed in individuals with SCI compared to able-bodied control groups using magnetic resonance imaging (MRI) (Henderson et al., 2011; Jurkiewicz et al., 2007; Karunakaran et al., 2019). Jurkiewicz and colleagues reported increased brain activation in the primary motor cortex as motor recovery improved after SCI, while activation in the associated sensorimotor regions was found to decrease (Jurkiewicz et al., 2007). Functional brain connectivity changes have also been observed in the thalamus and its subnuclei in individuals with complete paraplegic SCI (Karunakaran et al., 2020). Furthermore, MRI studies have shown that the brain’s gray matter volume is significantly atrophied in individuals with SCI compared to able-bodied controls (Karunakaran et al., 2019). These structural and functional brain changes may be associated with cognitive decline in individuals with SCI. Although both structural and functional MRI (fMRI) have been used to study brain reorganization after SCI, the rigid supine-positioning inside the scanner bore and magnetic contraindications pose limitations for investigating supraspinal changes after SCI, particularly in rehabilitative settings. In order to study brain reorganization after SCI more systematically or longitudinally, a more portable neuroimaging device such as functional near-infrared spectroscopy (fNIRS) may be used.

fNIRS is a non-invasive neuroimaging modality which utilizes near-infrared light to image the brain and quantify levels of hemoglobin concentration (Delpy et al., 1988; Jöbsis, 1977; Villringer et al., 1993). Near-infrared light is emitted through an optode source and collected by photodiode detectors, arranged across the brain depending on the regions of interest. Similar to fMRI, fNIRS measures neurovascular activity, based on the theory of neurovascular coupling, in which increased neural activity is followed by increased regional cerebral blood flow to a particular brain region (Girouard & Iadecola, 2006). Increased neuronal activation during an active task increases the demand for cerebral metabolites in specific task-associated regions, causing a cascade of changes in the cerebral blood flow, cerebral blood volume, oxygen metabolic rate, and consequently the oxygenated and deoxygenated hemoglobin concentration levels (HbO and HbR, respectively) (Buxton, 2010; Paulson et al., 2010). The use of fNIRS has become widespread in rehabilitation research due to its portability, high temporal resolution, robustness against motion artifacts, and ease-of-use (Yücel et al., 2017).

Despite the increasing use of fNIRS, it has not been widely used to study brain changes in individuals with SCI. We have previously used fNIRS to investigate functional brain changes in individuals with SCI compared to healthy controls and observed an overall decrease in functional brain hemodynamic response during finger tapping and ankle tapping tasks (Karunakaran et al., 2022). fNIRS has also been used in conjunction with robot-assisted gait training in individuals with SCI to measure functional brain activity changes in the primary motor cortex (Simis et al., 2018), in addition to being combined with repetitive transcranial magnetic stimulation to investigate the amelioration of neuropathic pain after SCI (Sun et al., 2019). These studies suggest that fNIRS can be a useful neuroimaging tool for investigating functional brain changes which occur in individuals with SCI. Not only can fNIRS be useful for investigating functional brain changes in the motor cortex regions, but it can also be used to investigate the cognitive deficits that have been reported in individuals with SCI (Alcántar-Garibay et al., 2022; Sachdeva et al., 2018). There is currently a lack of fNIRS studies investigating cognitive function in individuals with SCI. This is a much-needed area, particularly since studies investigating cognitive function in individuals with SCI have reported both significant cognitive impairments (Chiaravalloti et al., 2018; Craig et al., 2017; Davidoff et al., 1992; Dowler et al., 1997; Hall et al., 1999; Macciocchi et al., 2013; Molina et al., 2018; Wilmot et al., 1985) and no significant impairments (Bradbury et al., 2008; Masedo et al., 2005; Nott et al., 2014; Sachdeva et al., 2018). The conflicting reports of cognitive impairment in individuals with SCI could be partly attributable to the heterogeneity of the sample population, differences in the cognitive tests utilized, differences in study design, or sample size (Sachdeva et al., 2018; Sandalic, Tran, et al., 2022). Therefore, due to our currently limited understanding of cognitive impairment in individuals with SCI, fNIRS may be a useful tool to better understand cognitive impairment by quantifying concrete functional brain activity changes in individuals with SCI.

In the present study, we investigated the neurovascular correlates of working memory in individuals with SCI in comparison to able-bodied (AB) individuals using fNIRS and an N-back task. The N-back task is a frequently used task paradigm in fMRI and fNIRS studies to investigate working memory function and has been shown to activate frontoparietal regions of the brain (Braver et al., 1997; Owen et al., 2005). We also evaluated cognitive function in individuals with SCI and AB controls by using a brief cognitive screening tool established by Walterfang and colleagues, known as the Neuropsychiatric Unit Cognitive Assessment Tool (NUCOG) (Walterfang et al., 2006). Measures of brain hemodynamic activity were collected using fNIRS and correlated with accuracy, response time, and d-prime scores on the N-back task, in addition to NUCOG scores. The fNIRS-derived brain activity metrics, N-back performance scores, NUCOG scores, and brain-behavior associations were compared between the SCI and AB group.

## 2. Materials and Methods

### 2.1. Participants

19 individuals with SCI (14M, 5F; mean age ± standard deviation: 46.32 ± 10.18 years) and 25 age- and sex-matched able-bodied (AB) individuals (19M, 6F; mean age ± standard deviation: 43.2 ± 12.28 years) enrolled in the current study. All participants were recruited from the New Jersey/New York metropolitan area and provided written informed consent. Institutional Review Board (IRB) approval was obtained from New Jersey Institute of Technology. The exclusion criteria for the SCI group were as follows: (1) within one-year post-spinal cord injury or in the acute phase of injury, (2) presence of tetraplegia or inability to perform upper limb motor movements, (3) history of or concurrent traumatic brain injury (TBI), (4) history of or concurrently has psychiatric disorders such as post-traumatic stress disorder, addiction, bipolar disorder, or schizophrenia, (5) presence of acute illness or infection, (6) history of chronic hypertension, diabetes mellitus, stroke, epilepsy or seizure disorders, multiple sclerosis, Parkinson’s disease, (7) illicit drug abuse within the past 6 months, (8) has Alzheimer’s disease or dementia, (9) not able to speak English, and (10) younger than 18 years of age or older than 65 years of age. For the AB group, the same exclusion criteria were applied; however, individuals in this group did not have any presence or history of SCI. The inclusion criteria were as follows: (1) have spinal cord injury with more than one-year post-injury duration (SCI group only), (2) voluntary movement of the arms, (3) within 18-65 years of age, and (4) able to speak English. Injury characteristics such as the spinal level of injury, duration since injury, and completeness of injury were obtained from all individuals with SCI (Table 1).

**Table 1.**
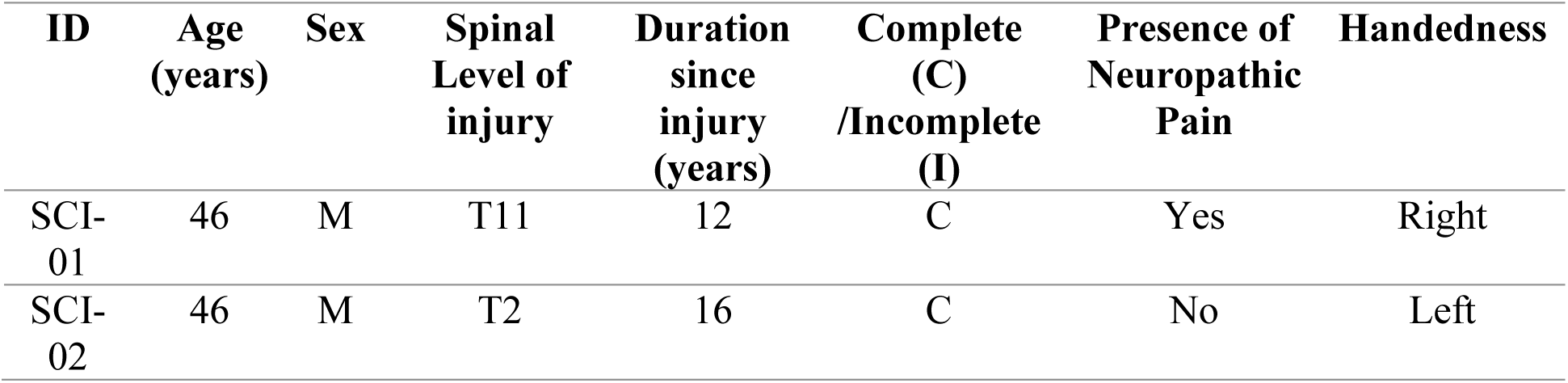
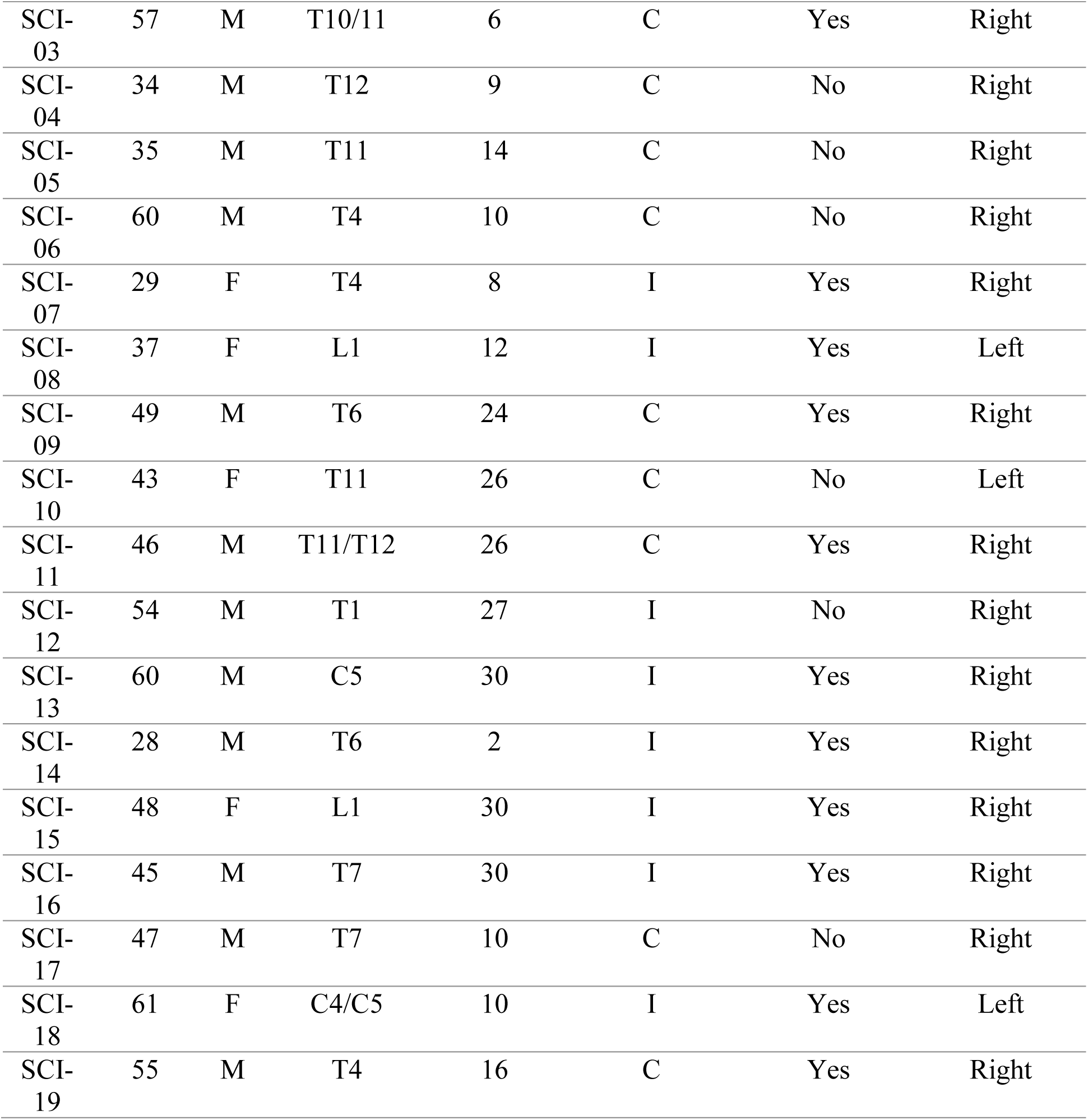
Demographic information and clinical characteristics from participants with SCI enrolled in the study.

| ID | Age<br>(years) | Sex | Spinal<br>Level of<br>injury | Duration<br>since<br>injury<br>(years) | Complete<br>(C)<br>/Incomplete<br>(I) | Presence of<br>Neuropathic<br>Pain | Handedness |
| --- | --- | --- | --- | --- | --- | --- | --- |
| SCI-01 | 46 | M | T11 | 12 | C | Yes | Right |
| SCI-02 | 46 | M | T2 | 16 | C | No | Left |
| SCI-03 | 57 | M | T10/11 | 6 | C | Yes | Right |
| SCI-04 | 34 | M | T12 | 9 | C | No | Right |
| SCI-05 | 35 | M | T11 | 14 | C | No | Right |
| SCI-06 | 60 | M | T4 | 10 | C | No | Right |
| SCI-07 | 29 | F | T4 | 8 | I | Yes | Right |
| SCI-08 | 37 | F | L1 | 12 | I | Yes | Left |
| SCI-09 | 49 | M | T6 | 24 | C | Yes | Right |
| SCI-10 | 43 | F | T11 | 26 | C | No | Left |
| SCI-11 | 46 | M | T11/T12 | 26 | C | Yes | Right |
| SCI-12 | 54 | M | T1 | 27 | I | No | Right |
| SCI-13 | 60 | M | C5 | 30 | I | Yes | Right |
| SCI-14 | 28 | M | T6 | 2 | I | Yes | Right |
| SCI-15 | 48 | F | L1 | 30 | I | Yes | Right |
| SCI-16 | 45 | M | T7 | 30 | I | Yes | Right |
| SCI-17 | 47 | M | T7 | 10 | C | No | Right |
| SCI-18 | 61 | F | C4/C5 | 10 | I | Yes | Left |
| SCI-19 | 55 | M | T4 | 16 | C | Yes | Right |

### 2.2. Cognitive Testing

All participants were administered the Neuropsychiatric Unit Cognitive Assessment Tool (NUCOG), which is a cognitive screening test that evaluates multiple cognitive domains such as attention, memory, visuoconstructional ability, executive function, and language skills (Walterfang et al., 2006). The duration of this test is approximately 20-30 minutes and scores for each of the sub-domains are calculated out of 20 points. The composite NUCOG score is calculated as the sum of the scores from all five cognitive domains. The NUCOG test has been evaluated on patients with neuropsychiatric disorders, showing sensitivity in differentiating patients with dementia and psychiatric subgroups, and has shown to be highly correlated with Mini-Mental State Examination (MMSE) scores (Walterfang et al., 2006). This test has also been used in evaluating cognitive impairment in individuals with SCI, showing lower scores overall in adults with SCI compared to AB adults across all five domains of the NUCOG test (Craig et al., 2016, 2017). For the purpose of this study, the NUCOG test was administered to provide a simple and streamlined method to evaluate potential cognitive differences between SCI and AB groups.

### 2.3. N-back Task

The N-back task was implemented with three separate levels of cognitive load: 0-back, 2-back, and 3-back, with stimuli presented via a computer screen, using E-Prime 2.0 software (Psychology Software Tools, Pittsburgh, PA) (Figure 1). The 1-back task was not used in this study due to its simplicity and similarity in cognitive load to the 0-back task. Participants were instructed to click either a left or right arrow key on a computer keyboard based on the letter stimulus displayed on the screen. With increasing *N*, the working memory load increased for participants, thus higher *N* is associated with greater task difficulty (Kane et al., 2007). Each N-back task consisted of three blocks, interleaved with 30 seconds of a resting period in which participants were instructed to not click any buttons. Within each block of the N-back task, participants were shown white letters on a black screen in succession, with an inter-trial interval of 2 seconds and 30 total letters, in pseudo-randomized order consistent across participants.

**Figure 1.**
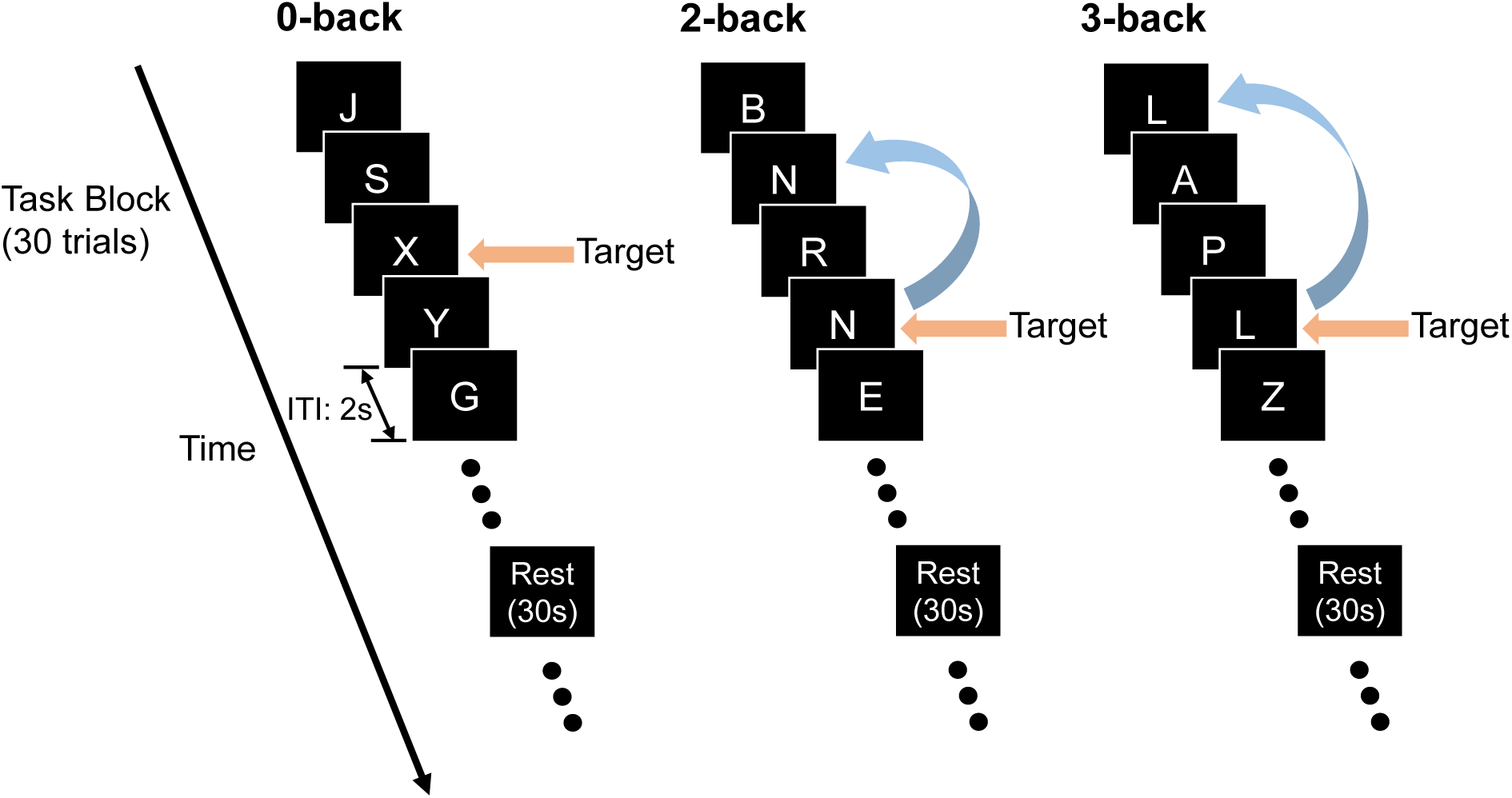
N-back task design showing each block and each task level, with examples of letters shown on a computer screen. ITI = inter-trial interval. For each N-back task level, a total of three blocks were implemented. Participants were asked to press a ‘right’ computer key on all target letters and press a ‘left’ computer key for all non-target letters.

Across all N-back tasks, the number of letters (or trials) were the same; however, different numbers of target and non-target letters were used based on the N-back task level. For the 0-back task, there were 10 target letters and 20 non-target letters for each block; for the 2-back task, there were 8 target and 22 non-target letters per block; and lastly, for the 3-back task, there were 7 target and 23 non-target letters per block. In the 0-back task, participants were instructed to click the right arrow key only when the target letter ‘X’ appeared on the screen; otherwise, participants were instructed to click the left arrow key. For the 2-back task, participants were instructed to click the right arrow key only if they saw the same letter two letters ago; otherwise, they were instructed to click the left arrow key. Lastly, for the 3-back task, participants were instructed to click the right arrow key on the target letter, defined as the same letter that appeared three letters ago (Figure 1).

### 2.4. fNIRS Data Acquisition

A continuous wave fNIRS system with 10 optode sources and 24 detectors was used, with 690 nm and 830 nm lasers (CW6 System, TechEn Inc., Milford, MA). A total of 30 channels were configured across the following regions of interest (ROI): right and left dorsolateral prefrontal cortex, medial prefrontal cortex, right and left inferior parietal lobe, and the right and left motor cortex (Figure 2). Four short source-detector separation (SDS) channels were configured at a distance of 0.84 cm between the source and detector, while 26 long SDS channels were placed at a distance of 3 cm between the source and detector. Short SDS channels are typically used to remove physiological signals from the extracerebral layers, recording information from the scalp and the cerebrospinal fluid (Brigadoi & Cooper, 2015). In the present study, data from the short SDS channels did not substantially improve the signal to noise ratio (SNR); thus, only data from the 26 long SDS channels were used and reported for subsequent analyses. Channel locations were determined and positioned with the aid of Brain Sight Neural Navigator (Rogue Research Inc. Neuronavigation System, Canada). For each participant, anatomical landmarks were collected, and the standard Montreal Neurological Institute (MNI) brain template was transformed into each individual’s space. All channels were placed according to MNI coordinates pre-defined in the literature (Table 2).

**Figure 2.**
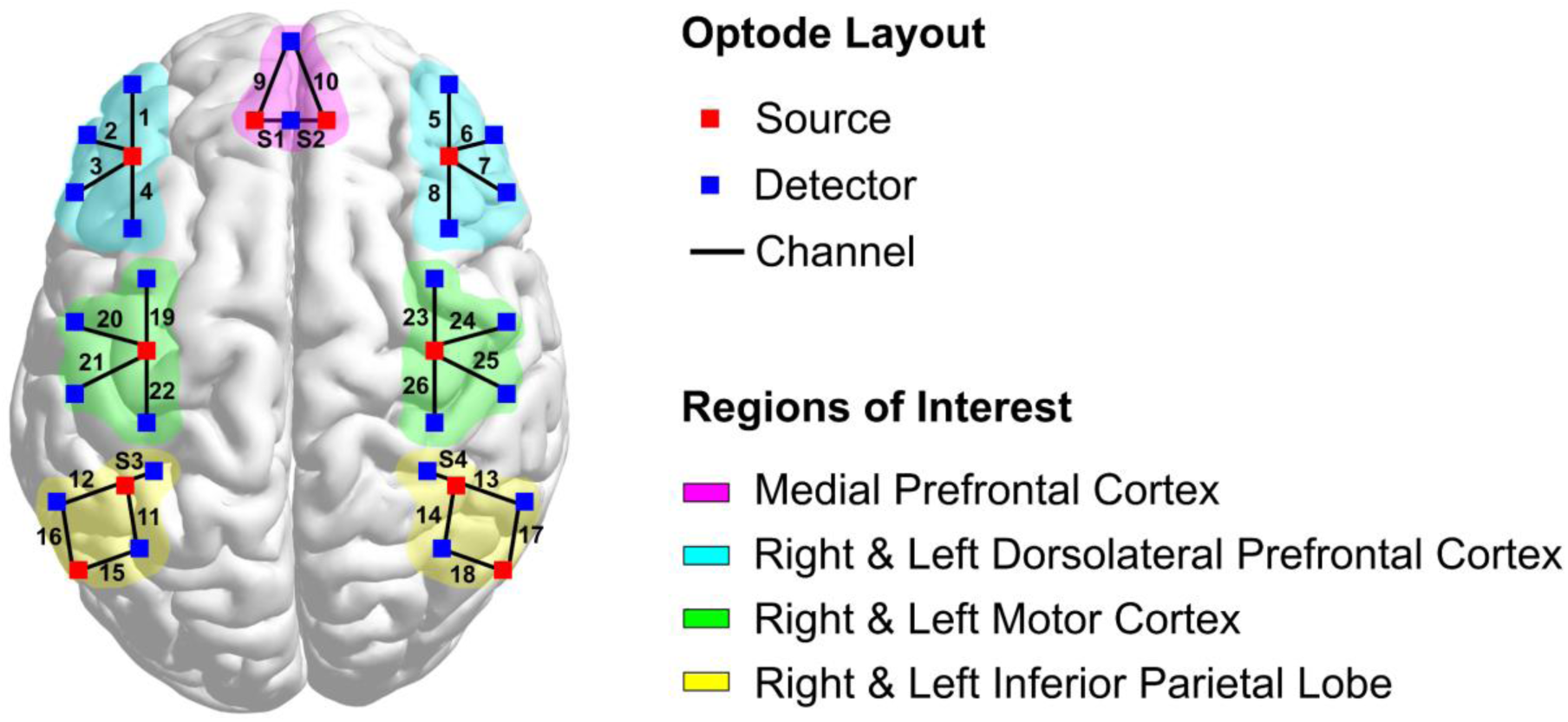
Optode layout design used for the fNIRS study. The channel #, source, and detectors are displayed, along with the following regions of interest: medial prefrontal cortex, right & left dorsolateral prefrontal cortex, right & left motor cortex, and right & left inferior parietal lobe. There are a total of 26 channels and 4 short source-detector separation channels (S1-S4), containing 10 sources and 24 detectors.

**Table 2.** Montreal Neurological Institute (MNI) coordinates displayed for each region of interest (ROI) used in the fNIRS setup. PFC: prefrontal cortex; DLPFC: dorsolateral prefrontal cortex.

| ROI | X | Y | Z | Reference | Channel #'s |
| --- | --- | --- | --- | --- | --- |
| Right DLPFC | 48 | 21 | 38 | (Watanabe et al., 2013) | 5, 6, 7, 8 |
| Left DLPFC | -48 | 21 | 38 | (Watanabe et al., 2013) | 1, 2, 3, 4 |
| Medial PFC | 0 | 61 | 22 | (Brier et al., 2012; Greicius et al., 2004;<br>Sestieri et<br>al., 2011) | 9, 10 |
| Right Motor<br>Cortex | 40 | -23 | 53 | (Brier et al.,<br>2012;<br>Ellermann et<br>al., 1998;<br>Fox et al.,<br>2009) | 23, 24, 25, 26 |
| Left Motor<br>Cortex | -40 | -23 | 53 | (Brier et al.,<br>2012;<br>Ellermann et<br>al., 1998;<br>Fox et al.,<br>2009) | 19, 20, 21, 22 |
| Right Inferior<br>Parietal Lobe | 52 | -54 | 36 | (Watanabe et<br>al., 2013) | 13, 14, 17, 18 |
| Left Inferior<br>Parietal Lobe | -52 | -54 | 36 | (Watanabe et<br>al., 2013) | 11, 12, 15, 16 |

### 2.5. Preprocessing

The raw fNIRS intensity data were first converted to optical density values, then wavelet-based head-motion correction was performed using 1.5 times the interquartile range as a threshold to remove outliers (Molavi & Dumont, 2012). Data were then bandpass filtered using a pass-band from 0.01 to 0.15 Hz to remove physiological noise sources such as cardiac rate and respiratory rate, while keeping the task frequency into consideration. The data were then converted from optical density values to oxygenated hemoglobin (HbO), deoxygenated hemoglobin (HbR), and total hemoglobin (HbT) values. Due to the highest signal to noise ratio observed in the HbO signal, subsequent analyses are shown for the HbO data. Data preprocessing was performed using custom scripts in MATLAB R2022b (The MathWorks Inc., Natick, Massachusetts) and HOMER2 functions (Huppert et al., 2009).

### 2.6. Behavioral Scores

To evaluate performance, accuracy and response time were collected for each block and each N-back task across all participants. Accuracy was calculated for each block of the N-back task and is defined as the number of correct responses divided by the total number of responses. Accuracy was calculated for all three block periods of the N-back task separately, as well as averaged together to obtain a mean accuracy score across all blocks for each participant. Furthermore, the response time was collected from each participant during each N-back task trial and the mean response time was obtained by averaging across all trials in each block. Sensitivity was also calculated to evaluate performance on the N-back task, due to the imbalance in number of target and non-target trials. Sensitivity was quantified by calculating d-prime scores, which were obtained by subtracting the z-transform of the false alarm rate from the z-transform of the hit rate (Stanislaw & Todorov, 1999). The hit rate was defined as the number of times the participant responded correctly to the target letters divided by the total # of targets, whereas the false alarm rate was defined as the number of times the participant responded incorrectly at non-target letters divided by the total number of non-target letters. In the case that the hit rate or false alarm rate values were equal to 1, the value was replaced by 1 – 1/(2*n), where n = total # of hits. Additionally, if the hit rate or false alarm rate values were equal to 0, then the value was replaced with 1/(2*n), according to Macmillan and Kaplan’s rule for handling 1’s and 0’s (Macmillan & Kaplan, 1985).

### 2.6. fNIRS Task Activation Metrics

For the N-back fNIRS data, block averaging and general linear model (GLM) analysis were performed. For block averaging, the time-series data were averaged for each block of the N-back task, as defined by 60 seconds of task followed by 30 seconds of rest. For each block, data collected during the 5 seconds prior to the start of each task stimulus were averaged and subtracted from the rest of the block’s time-series data, to normalize the data. Subsequently, each of the three blocks were averaged to generate a block-averaged hemodynamic response curve for each channel. Characteristics from the block-averaged data were extracted as task activation metrics, as follows: the area under the curve (AUC) of the hemodynamic response, time-to-peak (TTP), maximum HbO (maxHbO), and standard deviation of the HbO (stdHbO) signal. The maxHbO and stdHbO signals were obtained using the concatenated time-series data from all three blocks. For GLM, the N-back task was represented as a box-car function convolved with the canonical hemodynamic response function (Ashburner et al., 2014). This convolved hemodynamic response signal was input as the task regressor in the GLM model, with the preprocessed HbO time-series input as the outcome variable, and the beta coefficient was estimated for each channel to represent task activation. For each of the fNIRS metrics, the values were z-scored across all channels, for each participant. Each of the fNIRS task activation metrics were then correlated with performance metrics on the N-back task: accuracy, response time, and d-prime scores.

### 2.7. Statistical Analysis

All statistical analyses were performed using R 4.2.3 (R Core Team, 2023). For the N-back behavioral scores, a mixed-design ANOVA was performed for accuracy and response time values separately, with the following independent variables: group (SCI vs. AB), block (first, second, vs. third), and task (0-back, 2-back, vs. 3-back). The “block” and “task” variables were treated as within-subjects factors while the “group” variable was treated as a between-subjects factor. Post-hoc analyses were subsequently performed using paired t-tests when comparing between N-back task levels or block periods, while independent-samples t-tests were performed when comparing between the AB vs. SCI group. To correct for multiple comparisons, Bonferroni corrections were performed accordingly. For the fNIRS metrics, similar mixed-design ANOVAs were performed with each fNIRS metric constituting a separate independent variable and independent-samples t-tests were performed for each metric, task, and channel, with Bonferroni corrections performed for post-hoc multiple comparison testing. For all correlation analyses, Pearson’s correlation method was used, with Benjamini-Hochberg’s false discovery rate corrections applied for multiple testing across brain regions.

## 3. Results

### 3.1. NUCOG Scores

We observed no significant differences in the composite NUCOG scores between AB and SCI groups (independent-samples t-test, t(42) = 0.93, p = .36, 95% confidence interval: [-1.43, 3.87]), with the AB group showing a slightly higher mean NUCOG score than the SCI group (AB: 92.38 ± 4.50, SCI: 91.16 ± 4.05) (Figure 3A). Since the composite NUCOG score is composed of 5 distinct cognitive domains, these sub-domains were further analyzed. Significant differences were observed in the executive function sub-domain of the NUCOG test, in which the SCI group scored significantly lower than the AB group (Figure 3B) (independent-samples t-test, t(42) = 2.98, p = .0047, 95% confidence interval: [0.35, 1.81], Bonferroni adjusted α = .01 (.05/5). No significant differences between the two groups were observed across the other cognitive domains such as attention, visuoconstructional, memory, and language skills (Figure 3B).

**Figure 3.**
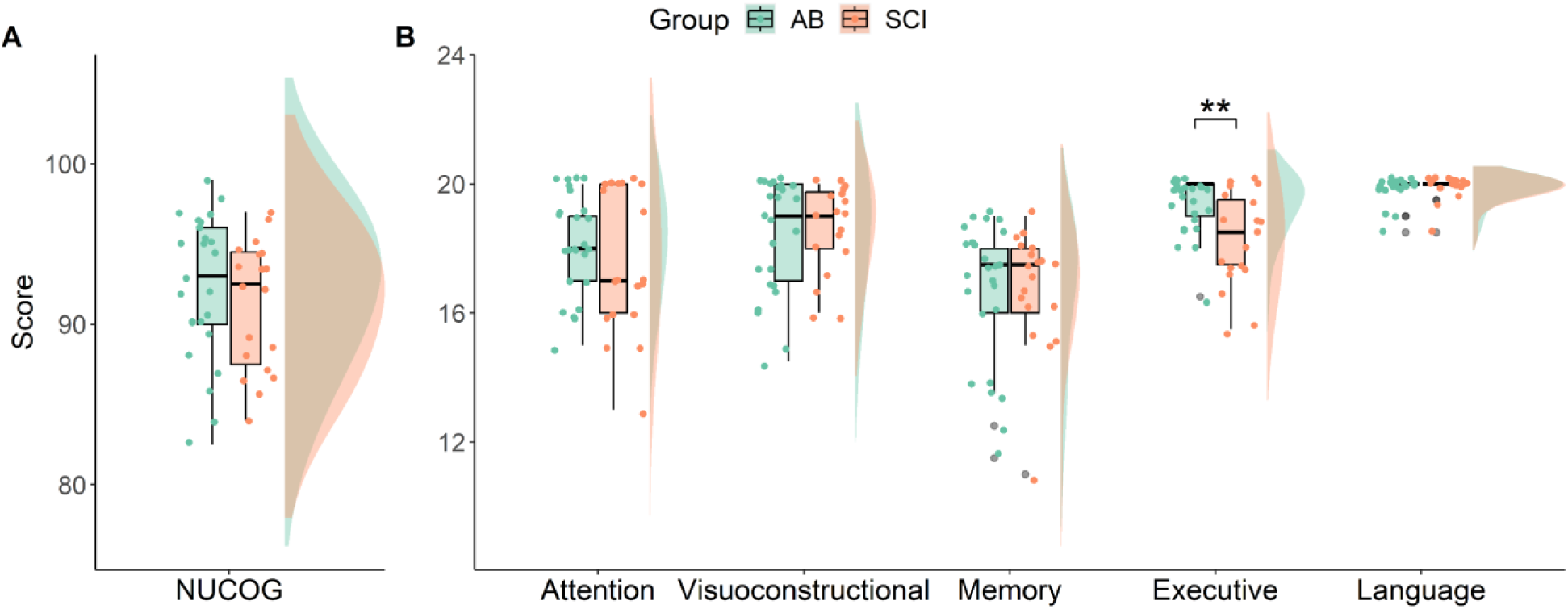
**A)** Neuropsychiatric Unit Cognitive Assessment Tool (NUCOG) scores for both able-bodied (AB) and spinal cord injury (SCI) groups are shown in green and orange bars, respectively. **B)** Sub-domains of the NUCOG test scores are also indicated, with significant differences shown between the AB and SCI group in executive function (independent-samples t-test, t(42) = 2.98, p = .0047, 95% confidence interval: [0.35, 1.81], Bonferroni adjusted α = .01 (.05/5)). The total scores from the NUCOG subdomains: attention, visuoconstructional, memory, executive, and language, are added to obtain the composite NUCOG score. ** p < .01.

### 3.2. N-back Task Performance

For all three N-back task behavioral scores: accuracy, response time, and d-prime, significant differences were observed between the N-back tasks and between blocks of the 2-back task in particular (Figure 4A-C). We found no significant differences between the SCI and AB groups for accuracy (mixed-design ANOVA, F(1,42) = 0.601, p = .44), response time (mixed-design ANOVA, F(1,42) = 0.188, p = .67), and d-prime scores (mixed-design ANOVA, F(1,42) = 1.76, p = .19). Across all behavioral metrics, there were significant effects of the N-back task level (0-back, 2-back, and 3-back levels) (mixed-design ANOVA; accuracy: F(2,84) = 72.43, p = 5.22e-19; response time: F(2,84) = 81.94, p = 1.83e-20; d-prime: F(2,84), p = 2.82e-07). This shows that the N-back task level contributes significantly to the behavioral scores. Within each task level, there were three blocks, and significant block effects were found for all three behavioral metrics (mixed-design ANOVA; accuracy: F(2,84) = 3.15, p = .048; response time: F(2,84) = 43.24, p = 1.23e-13; d-prime: F(2,84) = 6.78, p = .002), in addition to an interaction effect between task level and block (mixed-design ANOVA; accuracy: F(4,168) = 6.30, p = 9.57e-05; response time: F(4,168) = 24.70, p = 4.28e-16; d-prime: F(4,168) = 6.35, p = 8.81e-05). Post-hoc paired t-tests revealed significant differences in accuracy scores between the 0-back and 3-back tasks, as well as the 2-back and 3-back tasks for both SCI and AB groups, across all blocks (Bonferroni corrected α = .0027 (.05/18), all p-values < .0027) (Figure 4A). For response time, across all blocks, significant differences were found between the 0-back and 2-back task, and between the 0-back and 3-back task (Bonferroni corrected α = .0027 (.05/18), all p-values < .0027), but no significant differences were observed between the 2-back and 3-back tasks (Figure 4B). Lastly, d-prime scores revealed significant differences between the 0-back and 2-back task only, across all blocks (Bonferroni corrected α = .0027 (.05/18), all p-values < .0027) (Figure 4C).

**Figure 4.**
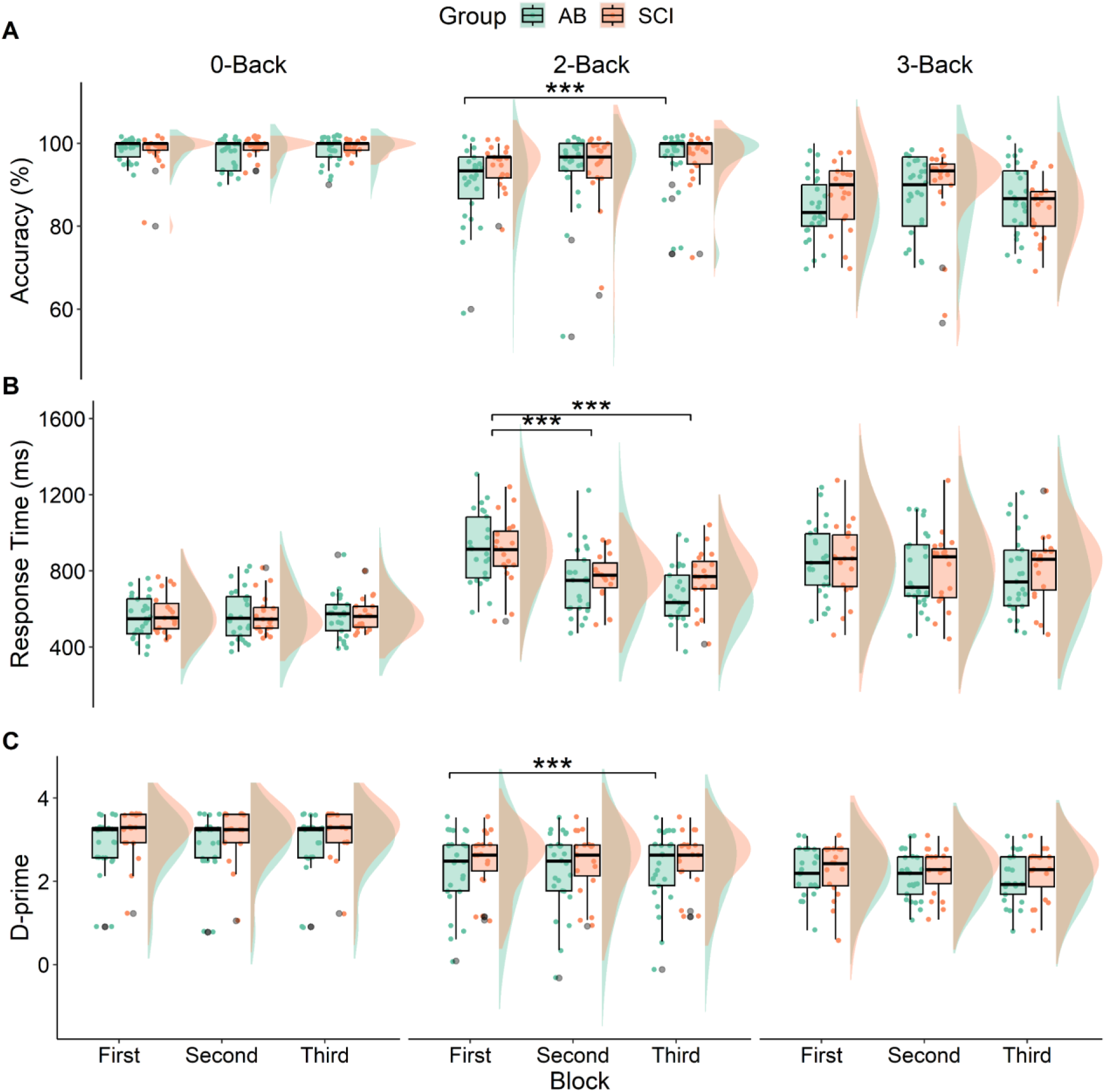
**A)** Accuracy, **B)** response time, and **C)** d-prime scores are shown for both spinal cord injury (SCI) and able-bodied (AB) groups, for each N-back task (0-Back, 2-Back, and 3-Back) and each block (first, second, and third). *** p < .001.

Between blocks, we found that performance was not constant and showed significant differences particularly during performance of the 2-back task (Figure 4A-C). For accuracy on the 2-Back task, there were significant differences in the AB group, between the first block and second block (paired-samples t-test with Bonferroni adjusted α = .0027 (.05/18), p = 6.83e-04) (Figure 4A). For response time on the 2-Back task, there are significant differences in the SCI group, between the first block and second block (paired-samples t-test with Bonferroni adjusted α = 0.0027 (.05/18), p = 9.03e-05), and between the first block and third block (paired-samples t-test with Bonferroni adjusted α = .0027 (.05/18), p = 1.09e-04) (Figure 4B). Similarly, significant differences for response time on the 2-Back task were observed for the AB group as well, between the first block and second block (paired-samples t-test with Bonferroni adjusted α = .0027 (.05/18), p = 1.28e-09), and between the first block and third block (paired-samples t-test with Bonferroni adjusted α = .0027 (.05/18), p = 1.93e-08) (Figure 4B). For d-prime scores, significant differences were observed between the second block and third block, only for the AB group (paired samples t-test with Bonferroni adjusted α = .0027 (.05/18), p = .0015) (Figure 4C).

### 3.3. fNIRS Task Activation Differences

The SCI group had a significantly higher maxHbO value than the AB group in channel #14, which corresponds to the right inferior parietal lobe, during the performance of the 2-back task (independent-samples t-test, p = .0018, FDR-correction at α=0.05, adjusted p = .0473) (Figure 5B&E. In the same region, the AUC of the hemodynamic response curve was also higher in the SCI group compared to the AB control group; however, results were not significant after false discovery rate (FDR) correction (independent-samples t-test, p = .044, FDR-correction at α=0.05, adjusted p = .57). Differences in task activation in channel #14 seem to be specific to the 2-back task and were not observed for the 0-back or 3-back tasks. During the performance of the 2-back task, there were some regions that showed higher task activation values in the AB group than SCI group, such as channels 19, 21, and 22 (Figure 5B&E), which are located in the left motor cortex. In channels 19 and 21, beta values were higher in the AB group than the SCI group, and in channel 22, the AUC, TTP, and maxHbO values were all higher in the AB group than the SCI group; however, the results were not significant after FDR correction. Most differences in task activation metrics between the AB and SCI groups were found in the 2-back task, while the 0-back task showed the least number of differences, being of lower cognitive load (Figure 5A&D). For the 3-back task, the SCI group showed higher beta values than that of the AB group, in channels 3 and 8 (left and right DLPFC, respectively); however, the results did not survive FDR correction (independent-samples t-test; channel 3: p = .022, adjusted p = .47; channel 8: p = .036, adjusted p = .47; FDR-correction at α=0.05) (Figure 5C&F).

**Figure 5.**
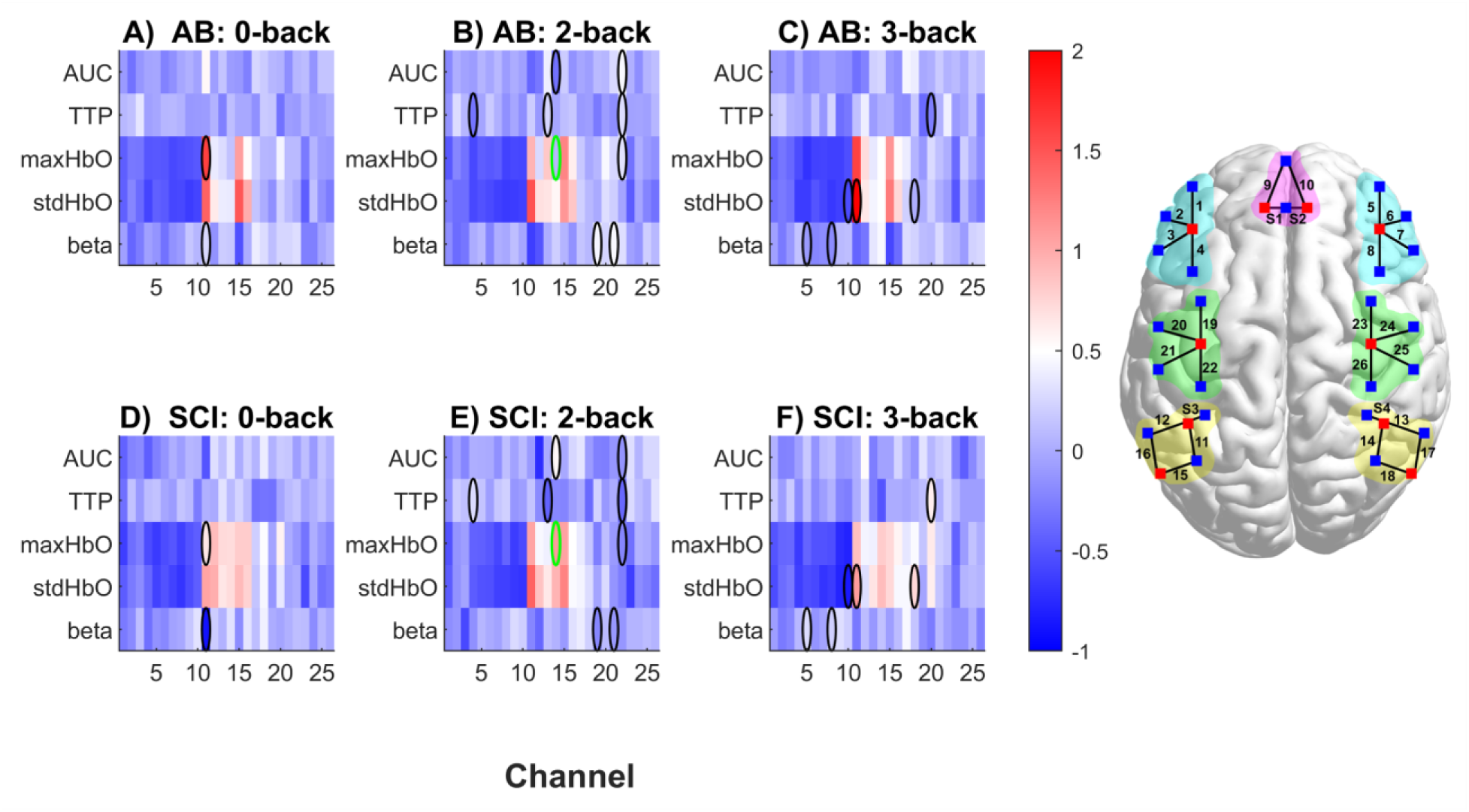
N-back working memory task fNIRS metrics between SCI and AB groups, across all 26 channels. The following fNIRS metrics (z-scored) are shown: area under the curve (AUC), time-to-peak (TTP), maximum HbO (maxHbO), standard deviation of HbO (stdHbO), and beta values from GLM analysis. Black ovals indicate significant differences between AB and SCI groups before correcting for multiple comparisons (p<.05). The green oval in maxHbO, Channel #14, during the 2-back task shows significant differences between the AB and SCI groups after FDR correction, α = 0.05. FDR correction was performed for each N-back task and each fNIRS metric, across all channels.

The overall task activation patterns across channels were similar between the SCI and AB groups, such as overall increased activation in maxHbO and stdHbO from channels 11-15, which correspond to areas of the inferior parietal lobes, as compared to the other channels (Figure 5A-F). This pattern was apparent across all levels of the N-back task. There were significant effects of the channel # on the following metrics: maxHbO, stdHbO, and beta values (mixed-design ANOVA; maxHbO: F(25, 1000) = 16.73, p = 1.00e-59; stdHbO: F(25, 1000) = 23.16, p = 6.08e-82; beta: F(25, 1000) = 1.76, p = .012).

### 3.4. Task Activation and Behavioral Scores

Significant associations between fNIRS task activation metrics and performance on the N-back tasks were observed for both AB and SCI groups across all fNIRS metrics except for stdHbO, and across both accuracy and response time values, in different brain regions. These significant associations were only observed for the higher cognitive load N-back tasks such as the 2-back and 3-back tasks, with the majority found in the 3-back task. No significant relationship between fNIRS metrics and performance metrics were observed for the 0-back task. All correlations were corrected for multiple comparisons using FDR correction at α = 0.05. The following regions showed significant brain-behavior associations: left inferior parietal lobe (channel #12), right inferior parietal lobe (channel #18), and the right dorsolateral prefrontal cortex (channel #6 & #8) (Table 3). Furthermore, significant correlations between fNIRS task activation metrics and NUCOG scores were also observed, for the composite NUCOG scores, visuoconstructional, memory, and attention scores (Table 3).

**Table 3.** Pearson’s correlations are shown between fNIRS metrics and performance metrics that are significant after multiple comparison correction using the Benjamini-Hochberg’s false discovery rate (BH-FDR) method. Prior to correlating the metrics, outliers were removed in either variable based on the 1.5*IQR rule. TTP: time to peak; AUC: area under the curve; maxHbO: maximum oxy-hemoglobin concentration.

| Group | Task | Channel # | Performance metric | fNIRS metric | r-value | p-value | FDR adjusted p-value |
| --- | --- | --- | --- | --- | --- | --- | --- |
| **AB** | 2-Back | 12 | Accuracy | beta | -0.71 | 0.00091 | 0.024 |
| **AB** | 3-Back | 6 | Response Time | TTP | -0.65 | 0.00052 | 0.014 |
| **AB** | 3-Back | 18 | Accuracy | TTP | 0.63 | 0.0010 | 0.027 |
| **SCI** | 3-Back | 8 | Response Time | AUC | 0.71 | 0.00097 | 0.025 |
| **SCI** | 3-Back | 8 | Response Time | maxHbO | 0.75 | 0.00047 | 0.012 |
| **AB** | 2-back | 23 | NUCOG | AUC | 0.65 | 0.0013 | 0.035 |
| **AB** | 2-back | 23 | NUCOG | TTP | -0.60 | 0.0019 | 0.049 |
| **AB** | 3-back | 22 | NUCOG | TTP | 0.61 | 0.0014 | 0.037 |
| **AB** | 2-back | 8 | Visuo-constructional | beta | 0.65 | 0.0014 | 0.035 |
| **AB** | 2-back | 23 | Memory | AUC | 0.66 | 0.0013 | 0.033 |
| **SCI** | 2-back | 13 | Attention | AUC | 0.67 | 0.0016 | 0.042 |
| **SCI** | 3-back | 3 | Visuo-constructional | TTP | 0.69 | 0.0011 | 0.028 |
| **SCI** | 3-back | 26 | Visuo-constructional | beta | 0.70 | 0.0016 | 0.042 |

## 4. Discussion

In the present study, we investigated differences in cognitive function between individuals with SCI and AB controls using the NUCOG test and N-back task with three different cognitive loads of increasing difficulty. We also investigated differences in functional brain hemodynamic activity between the SCI and AB groups using fNIRS, and evaluated the relationship between the brain’s hemodynamic characteristics and cognitive performance metrics. On the NUCOG test, we observed significant differences between the SCI and AB groups on measures of executive function, but no significant differences were observed in other sub-domains of the NUCOG test. For the N-back task, across the three levels of difficulty, no significant differences were observed between the SCI and AB group; however, both groups performed worse as the level of difficulty increased. Although there were no significant differences in N-back performance scores between the two groups, functional brain hemodynamic activity differences were observed between the SCI and AB groups, particularly in the right inferior parietal lobe.

### 4.1. Lower Executive Function Scores in Individuals with SCI

We found no significant differences in performance on the composite NUCOG scores between individuals with SCI and AB controls; however, significant differences were observed in the executive function sub-domain of the NUCOG test. The executive function test contains questions involving motor sequencing, categorical fluency, abstract thinking, and managing interference (Walterfang et al., 2006). Lower scores in executive function could potentially be due to the motor impairment observed in individuals with SCI, since the executive function test contains a motor sequencing task, which relies on the participants following a 3-step sequence of hand movements, repeated 5 times with each hand. Although all participants had retained upper-body movement, some may have struggled with fine motor control of the hands, such as an SCI subject with C4-5 injuries, thus impacting the executive function score. Craig and colleagues observed reduced executive function in individuals with SCI compared to AB controls as well; however, they also reported reduced cognitive functions in the other NUCOG sub-domains as well, which were not apparent in our cohort (Craig et al., 2017). Furthermore, when administered the Delis-Kaplan Executive Function test for verbal fluency, individuals with SCI were shown to score significantly lower than age-matched healthy controls (Chiaravalloti et al., 2018). Cohen and colleagues evaluated executive function using flanker and card sorting tests, and also found that individuals with SCI performed significantly worse than controls, suggesting that cognitive screening tests for individuals with SCI may want to focus on targeting executive function (Cohen et al., 2017). In a meta-analysis conducted by Sandalic and colleagues, adults with SCI were found to experience deficits in cognitive functioning when compared to AB individuals, mostly in attention and executive functioning domains (Sandalic, Craig, et al., 2022). These studies are consistent with our findings of lower executive function scores in individuals with SCI as compared to AB controls. It is important to note that in evaluating the structural validity of the NUCOG test using congeneric confirmatory factor analyses, Sandalic and colleagues found that the total composite NUCOG score was sensitive to differences in neurocognitive capacity; however, subdomains such as executive function had poor fit, while attention and visuoconstruction had adequate model fit (Sandalic, Tran, et al., 2022).

### 4.2. Similar N-back Scores between SCI and AB groups

To our knowledge, this is the first study which has implemented the N-back task to study working memory function in individuals with SCI. The SCI and AB groups performed similarly on the N-back tasks throughout all three levels of difficulty. This finding is consistent with results from the NUCOG test, particularly in the memory subdomain, which also revealed no significant differences between the SCI and AB groups. We observed decreased performance in both groups with increasing N-back task level, which was expected, since increasing working memory load increases the difficulty of the task, resulting in decreased accuracy, longer response times, and decreased d-prime values (Lamichhane et al., 2020). Both groups performed the worst in the 3-back task and we expect that increasing the working memory load further would display a similar downwards trajectory in performance.

Within the 2-back task, we observed significant differences between blocks. Both SCI and AB groups had a significantly longer response time in the first block compared to the second and third blocks. This may be due to learning effects which took place, since the 2-back task is much more difficult than the 0-back task; therefore, there is more room for improvement in scores throughout the blocks. Improvements in N-back task performance have been observed both within and between-sessions (Yeung & Han, 2023), and has been used in paradigms to train working memory (Miró-Padilla et al., 2019; Soveri et al., 2017). It is important to note that improvements in N-back task performance may not necessarily transfer to other domains or other measures of cognitive abilities in healthy adults (Thompson et al., 2013); however, it may be useful for cognitive training for individuals with SCI with working memory impairments. Therefore, future studies may want to investigate the efficacy of the N-back task in training individuals with SCI with cognitive impairments.

### 4.3. Increased Brain Activation in Individuals with SCI during the 2-Back Task

Although we did not observe significant differences in N-back task performance metrics between individuals with SCI and AB controls, we found significant fNIRS task activation differences between the two groups. Significant differences in maximum hemoglobin concentration were observed in the right inferior parietal lobe during performance of the 2-back task. In this region, the SCI group had a higher maximum hemoglobin concentration value than the AB group, which may represent increased effort in performance of the 2-back task. The right inferior parietal lobe is consistently activated across subjects in n-back studies (Owen et al., 2005), and is involved in multiple functions, spanning from attention to action processing (Caspers et al., 2013). Brain activity has been found to be higher in more cognitively demanding tasks such as the 2-back task, as compared to lower cognitive load tasks such as the 0-back or 1-back task (Braver et al., 1997; Herff et al., 2014; Meidenbauer et al., 2021). We found greater similarity between the SCI and AB groups in the 0-back task, perhaps due to it being an easier task and involving fewer working memory faculties than the 2-back or 3-back tasks. Although the performance between the AB and SCI groups were not significantly different in the 2-back task, it is possible that the SCI group activated the right inferior parietal lobe more in order to focus and apply more cognitive effort to the task. Individuals who are trained with the N-back task over time typically show decreased activation in various brain regions including the inferior parietal cortex, particularly for higher load N-back tasks (Miró-Padilla et al., 2019; Vermeij et al., 2017; Yeung & Han, 2023). This likely resembles the lower amount of effort in task activation needed as individuals improve on the N-back task. Therefore, further cognitive training in individuals with SCI may reduce task activation in the right inferior parietal lobe and warrants further investigation.

### 4.4. fNIRS Task Activation and its Association with Behavioral Scores

We observed significant associations between fNIRS task activation metrics and performance scores on the N-back task, as well as on the NUCOG scores. The significant brain-behavior associations were observed for both SCI and AB groups; however, those in the SCI and AB groups showed significant associations in different regions of the brain. For the SCI group, increased response time in channel 8, corresponding to the right DLPFC, was associated with increased fNIRS task activation metrics, such as maxHbO and AUC values. Whereas for the AB group, significant associations were observed in channels 12, 6, and 18, in regions of the left IPL, right DLPFC, and right IPL regions, respectively. It is possible that brain-behavior interactions between the AB and SCI groups are differently expressed. No significant brain-behavior associations were observed for the 0-back task, which is consistent with Lamichhane and colleagues’ study on cognitive load in the N-back task, as they revealed that brain-behavior relationships are stronger in high load conditions (up to the 6-back task), as compared to low-load conditions, such as the 0-back and 1-back task (Lamichhane et al., 2020).

### 4.5. Limitations

It is important to note limitations of the current study, such as the limited number of fNIRS channels covering the brain. Having more channels covering would allow for greater spatial resolution and analyses on the interactions between distinct brain regions. Additionally, having a larger sample size would allow for further analysis of high-performers vs. low-performers, which may be investigated in future studies, particularly since there may be heterogeneity within the SCI group. The lower sample sizes may underpower some of these comparisons. In the current study, we only used the N-back task to evaluate working memory during fNIRS acquisition; however, it may be beneficial to also include other cognitive tasks such as the Stroop task and Flanker task to confirm if the differences observed in the N-back task are similar to that of other cognitive tasks.

## 5. Conclusion

Using fNIRS coupled with an N-back working memory task, the current study investigated differences in cognitive function between individuals with SCI and AB controls. We found increased activity in the right inferior parietal lobe in individuals with SCI during the 2-back task as compared to the AB group. However, no differences in N-back task performance were observed between the SCI and AB groups, despite differences in executive function scores on the NUCOG subdomain test.

## Acknowledgments

The authors would like to sincerely thank Dr. Christopher Cirnigliaro, Ms. Annie Kutlik, and Mr. Steven Knezevic from the Spinal Cord Damage Research Center, James J. Peters Veterans Affairs Medical Center, Bronx, NY for helping with patient recruitment for this study. Research reported in this publication was supported by the New Jersey Commission on Spinal Cord Research under Award Number CSCR22ERG026 to B.B.B. This work was also supported by the National Center for Advancing Translational Sciences (NCATS) of the National Institutes of Health (NIH) under Award Number TL1TR003019 to D.Y.C. The content is solely the responsibility of the authors and does not necessarily represent the official views of the funding agencies.

## Notes

### Competing Interest Statement

The authors have declared no competing interest.

